# Determination of the water network surrounding the type I pilus from *E. coli* by cryo-electron microscopy

**DOI:** 10.1101/2025.09.07.674696

**Authors:** T.E. Petrova, A.S. Glukhov, A. Stetsenko, A. Guskov, A.G. Gabdulkhakov

**Affiliations:** Institute of Mathematical Problems of Biology, Keldysh Institute of Applied Mathematics, RAS, Professor Vitkevich St., 1, Pushchino, 142290 Russian Federation; Institute of Protein Research, RAS, Institutskaja str, 4, Pushchino, 142290 Russian Federation; Groningen Biomolecular Sciences and Biotechnology Institute, University of Groningen, Groningen 9747AG, the Netherlands

**Keywords:** Type 1 pili from *E. coli*, cryo-EM reconstruction, water network

## Abstract

Type 1 pili are protein filamentous surface structures of Gram-negative bacteria that mediate adhesion to host and play a crucial role in infection. Here, we report the cryogenic electron microscopy structure of the type 1 pilus from *E. coli* K-12 comprising 15 subunits of the major protein pilin FimA. The final resolution of EM reconstruction was estimated to be in the range from 2.09 to 2.30 Å, which is higher than that of the previously published structure. This improvement in the resolution enabled us to refine side-chain conformations to reliably determine the distances between the side-chain residues participating in the intersubunit interactions, and determine a network of water molecules surrounding the pilus rod. The analysis revealed that water contributes to intersubunit stabilization both through discrete bridging interactions and through extended hydrogen-bonded clusters, thereby supporting both the rigidity and flexibility of the filament. Comparison with a homologous high-resolution pilus model from enterotoxigenic *E. coli* showed that nearly all “conserved” water molecules i.e., those that are present at equivalent positions in different subunits of our model occupy also equivalent positions across the two structures, underscoring their functional relevance. At the same time, sequence-specific differences in hydration patterns were observed. These findings highlight the structural and functional importance of water in pilus architecture and provide a more detailed molecular framework for understanding bacterial adhesion.

**Synopsis:** The improvement in the resolution of the Cryo-EM reconstruction for type I pilus from E. coli made it possible to determine the positions of water molecules surrounding the pilus rod and reveal a more detailed picture of interactions between different subunits of the rod.

## 1. Introduction

Type 1 pili are protein filamentous hair-like appendages on the surface of Gram-negative bacteria, which enable adhesion to host cells and thereby initiate infectious process [1,2]. In uropathogenic bacterium *Escherichia coli* (UPEC), pili-mediated attachment to epithelial cells of the urinary tract is essential for its virulence, making these structures important targets for understanding bacterial pathogenicity.

Type 1 pilus is primarily composed of the major pilin protein FimA, which has an immunoglobulin-like fold. FimA polymerizes into a long helical rod structure containing up to 3000 subunits and possessing helical symmetry, with ∼ 3.2 subunits per turn, stabilized by extensive intersubunit contacts [3,4]. The assembly occurs via the so-called “donor-strand exchange” mechanism, in which the N-terminal strand of each subunit is inserted into the incomplete fold of the preceding subunit [5]. Numerous interactions are found between subunits that are located on the neighboring turns of the helix. This architecture provides both remarkable stability and mechanical flexibility, enabling pili to withstand shear forces while maintaining adhesive function [6,7].

In addition to FimA, the pilus contains the adhesin FimH at its tip, the adaptor proteins FimF and FimG, which connect the rod to FimH, and several accessory proteins FimI, FimC, and FimD, required for assembly and structure integrity [8-10].

High-resolution structural information on FimA and type I pili has been obtained using X-ray crystallography, NMR, and cryo-electron microscopy (cryo-EM). Crystallographic studies of isolated FimA subunits have achieved resolutions as high as 0.89 Å for FimA from *S*.*flexnery* (PDB: 5lp9, [11]) and 1.5 Å for FimA from UPEC (PDB: 5nkt, [11]), providing atomic details of its fold. However, due to the challenges in crystallizing long filamentous assemblies, cryo-EM has become the method of choice for determining structures of pili and revealing the overall architecture and intersubunit organization. [4, 12]. Recent cryo-EM studies reported reconstructions of pili from UPEC at 2.52–2.85 Å [12].

Advances in detector technology [14] and image processing algorithms [15] now enable reconstructions at resolutions approaching 2 Å. Such improvements allow precise modeling of amino acid side chains and the visualization of individual water molecules. Since water often mediates protein–protein interactions, contributes to stability, and influences conformational flexibility, mapping the hydration network in pili is critical for understanding their mechanical properties and function.

In this study we present a cryo-EM reconstruction of type I pili from UPEC K-12 with improved resolution (2.09–2.30 Å). The higher resolution allowed us to improve the model and determine a comprehensive water network within the pilus rod, and to assess its structural and functional roles. The analysis of the water network surrounding the pilus rod is of great interest for analyzing the contacts between different subunits, to understand the role of water molecules in the flexibility of the pilus rod, and to investigate the possible involvement of water in the polymerization process and rod function.

During the manuscript preparation, the similar study has been published, in which water networks were determined from two high-resolution EM pili models, one of which was the model of type I pili from enteroxigenic *E. coli* (PDB: 9IUG, [16]). This allowed us to further validate our model by comparison with this homologous high-resolution pilus structure, revealing conserved water positions as well as sequence-dependent hydration differences. Our results provide new insights into the molecular determinants of pilus stability and flexibility, highlighting the importance of water in bacterial adhesion mechanisms.

## 2. Materials and methods

### 2.1. Purification of FimA

*E. coli* cells of the strain XAC were grown at 37°C in LB medium (10 g/L tryptone, 5 g/L yeast extract, 10 g/L NaCl) until the middle of the log growth phase. The cells were concentrated 10-fold by centrifugation (6,000g for 20 min) and subsequent dissolution in a buffer solution containing 50 mM Tris-HCl, pH 7.5, 200 mM NaCl, and 5 mM EDTA. Lysozyme was added to the cell suspension to a concentration of 1 mg/mL, and DNase I and RNase A were added to a concentration of 1 ng/ml. The suspension was incubated at 37°C for 1 h. Debris was precipitated by low-speed centrifugation at 6,000g for 40 min. PEG 6,000 and NaCl were added to the clarified lysate to a concentration of 9% and 0.5 M, respectively, and incubated at 4°C for 12 h. A sample was concentrated 40-fold by centrifugation at 6,000g for 20 min and subsequent dissolution in a buffer solution containing 50 mM Tris-HCl, pH 7.5, and 200 mM NaCl. The resulting solution was applied to a linear glycerol gradient (15-30%) and centrifuged at 24,000 rpm for 3 h (SW41Ti rotor). The linear gradient was then divided into 1 mL fractions and analyzed by 12% SDS-PAGE. The fractions containing the protein of interest were combined, dialyzed against a solution containing 50 mM Tris-HCl, pH 7.5, and 200 mM NaCl, and concentrated using an Amicon 100 ultrafiltration device (Millipore).

### 2.2. Cryo-EM grid preparation and data collection

The sample was concentrated with a buffer containing 50 mM Tris-HCl, pH 7.5, and 200 mM NaCl to the final concentration of 1.0 mg/mL. Then, 3 μL of the sample were applied onto freshly glow-discharged (30 s at 5 mA) Quantifoil grids (Au R1.2/1.3, 300 mesh) at 20°C and 100% humidity (blotting time 3-5s, blot force – 0) and plunged-frozen in liquid ethane. The cryo-EM data were collected using 300 keV Krios microscope (Thermo Fisher) at NeCEN facility, equipped with Gatan K3 detector and Bioquantum Energy Filter.

Statistics of the data collection, map reconstruction, and model refinement are summarized in Table 1.

**Table 1.** Cryo-EM data collection, refinement, and validation statistics.

|  |  |
| --- | --- |
| **Data collection** |  |
| Voltage (kV) | 300 |
| Magnification | 105,000 |
| Electron dose (e-/Å<sup>2</sup>) total | 50 |
| Defocus range (µm) | -0.5, -0.8, -1.1, -1.4, -1.7, -2.0 |
| Pixel size (Å) | 0.836 |
| No. of movies | 8018 |
| **Data processing** |  |
| Software | cryoSPARC |
| No. of final particles | 1,593,408 |
| Final map resolution | 2.09 |
| FSC threshold | 0.143 |
| Map sharpening B factor (Å<sup>2</sup>) | 77.4 |
| **Helical symmetry** |  |
| Point group | C1 |
| Helical rise (Å) | 7.738 |
| Helical twist (°) | 115.00 |
| **Refinement and Model validation** |  |
| Model composition |  |
| Atoms | 16197 |
| Protein residues | 2385 |
| Water molecules | 1644 |
| R.m.s.d. |  |
| Bond length (Å) | 0.007 |
| Bond angles (°) | 0.676 |
| Clashscore | 2.44 |
| Rotamer outliers (%) | 1.61 |
| Ramachandran plot (%) |  |
| Favored | 95.80 |
| Allowed | 4.20 |
| Disallowed | 0 |

### 2.3. Cryo-EM image processing and reconstruction

The collected cryo-EM movies were processed and analyzed using cryoSPARC [17]. Data processing was conducted using the embedded helical reconstruction functions. First, all movies were dose-weighted using the Patch Motion Correction procedure. Then, the contrast transfer function (CTF) parameters were estimated. 8018 CTF-corrected micrographs were selected for further processing. Filaments were automatically picked, and segments were extracted with a box size of 256px. The 2D classification was performed by skipping the first CTF peak and limiting the resolution to 3.6 Å. The best 2D classes were used as references for filament autopicking from the whole dataset. In total, approximately 20 M segments were extracted using the same box size of 256px, and the 2D classification was repeated using the same settings. The best-looking 2D classes comprising a total of 1767k particles were selected. The 2D class averages of the filament segments showed a homogeneous class amenable to performing the reconstruction at a resolution of 2.5 Å followed by the asymmetric helical refinement. The symmetric parameters were then estimated, and one clear solution, with a rise of 7.74 Å and a twist of 115.00° was found. Subsequent helical reconstruction with the use of these parameters yielded a high-quality cryo-EM density map.

The resolution of the reconstruction was estimated using the Fourier shell correlation (FSC) plot. For the raw unmasked map, the curve crosses the 0.143 threshold at a resolution of 2.5 Å, while the “tight” and “corrected” curves corresponding to masked maps intersect the 0.143 level at 2.09 Å (Fig. 1a). The global map sharpening with the uniform B-factor of 77.4 was applied.

**Figure 1.**
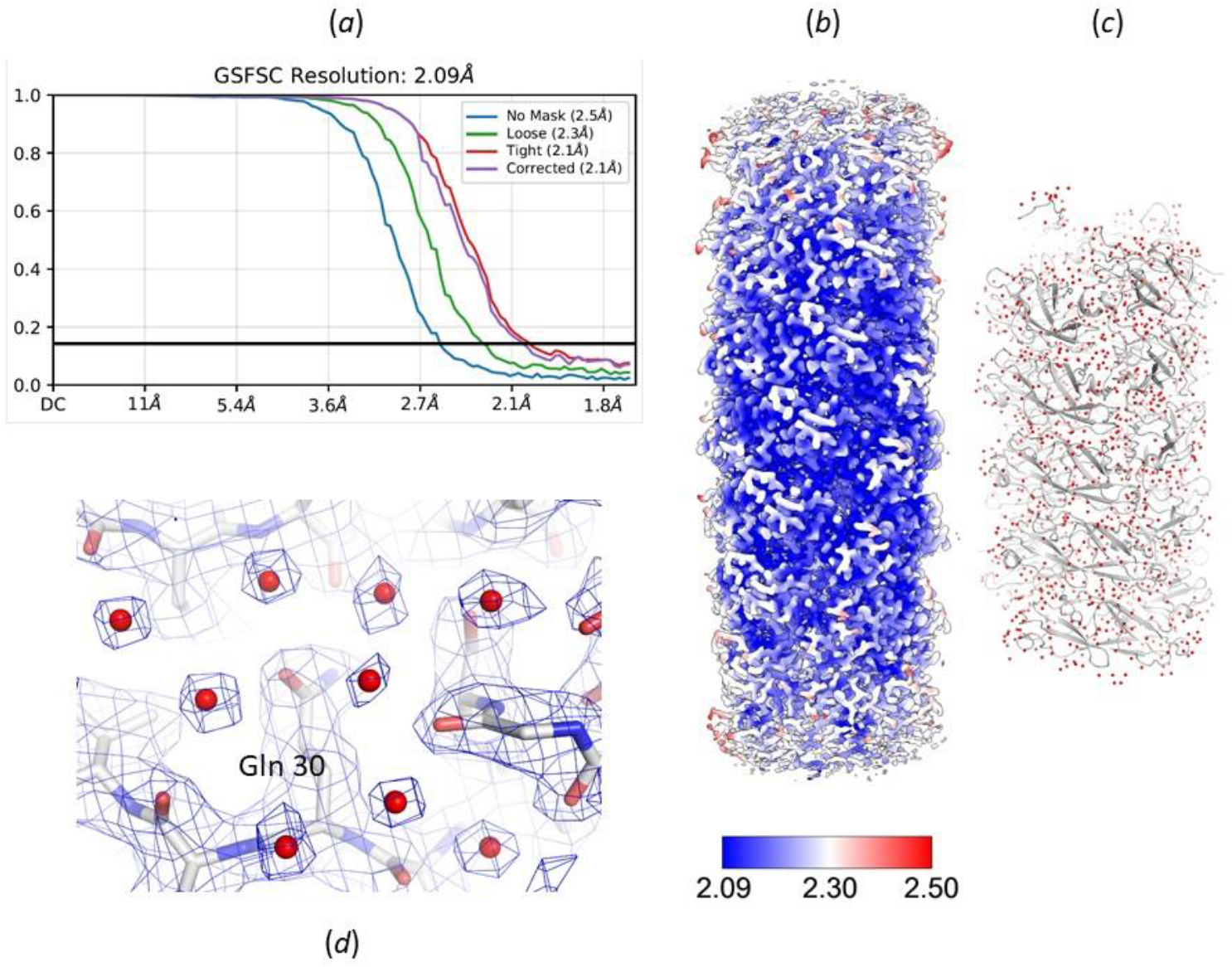
The EM reconstruction. (a) FSC-curve showing the correlation coefficient between successive spherical shells in Fourier space. The curve was calculated for the two half-maps. (b) The 3D surface view of the cryo EM-map at the contour level 0.204. The map is colored according to the local resolution value. (c) Cartoon representation of the pilus rod consisted of 15 subunits of the protein FimA and 1644 water molecules. (d) The water network on the inner surface of the pilus rod in the vicinity of Gln 30 of subunit G. The EM-map is shown at the contour level 0.22.

### 2.4. The building and refinement of the model

An initial model of a single subunit was constructed *de novo* without using the protein sequence. Then, model adjustment was performed using the models of the FimA protein of 16 kDa solved by X-ray diffraction at 1.5 Å resolution (PDB: 5nkt, [11]) and the cryo-EM models of type 1 pili rods (PDB: 6y7s and 8ptu, [13]). The models were fitted into the EM map using Chimera [18], and further model rebuilding was done in Coot [19]. The refinement of the model in direct space was performed using the phenix.real_space_refine module of the Phenix software package [20] initially against the unsharpened map followed by refinement against the sharpened map. Water molecules were added to the model using the phenix.douse procedure. The value of the map_threshold_scale parameters was chosen to be 0.3. Some water molecules were removed after manual inspection. The model quality check was performed using the MolProbity program [21]. The constructed pilus rod consists of 15 subunits (A - O), which is about 5 turns of the helix and 1644 water molecules. The map and the model are shown on Figs.1b-d and Fig.2.

**Figure 2.**
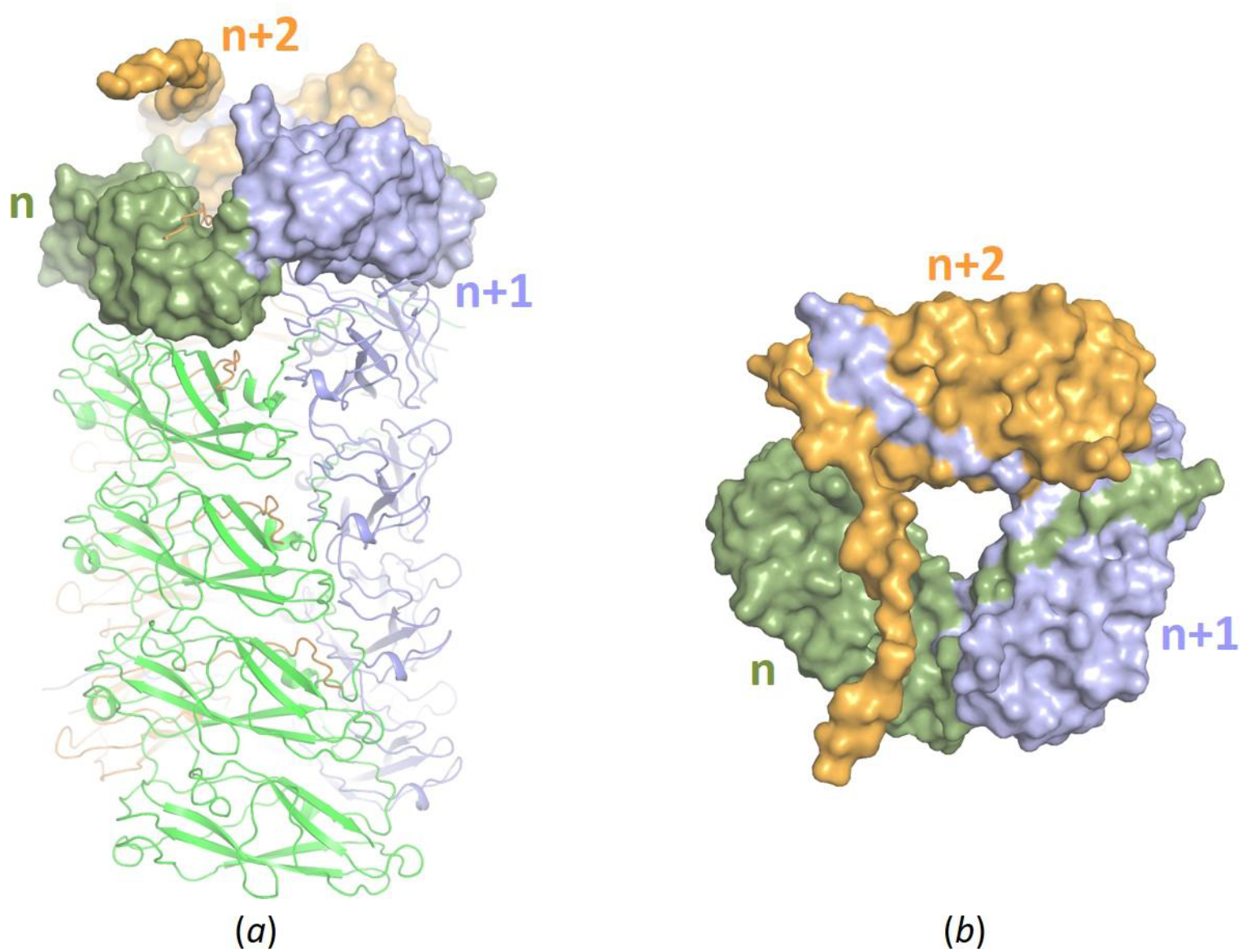
The helical assembly of the type I pilus from *E. coli* K-12. Each FimA subunit n has contacts with three subunits before and three subunits after n+1, n+2, and n+3 it along the course of the rod. Front (a) and top (b) views of the pilus rod.

Cryo-EM maps, both sharpened and unsharpened, have been deposited in the wwPDB OneDep System under the Electron Microscopy Data Bank accession code EMD-65144. The atomic model has been deposited in the PDB (PDB: 9VKV).

## 3. Results and discussion

### 3.1. Structure determination

The pilus rod model was refined in real space first against the raw unsharpened map and subsequently against a sharpened map generated by applying the uniform B-factor. Global B-factor map sharpening is the simplest and most widely used correction method in cryo-EM as it reduces the contrast loss at high resolution and enhances map interpretability [22]. Alternative strategies employ either global or local B-factor estimation for different map regions [23-25]. In our case, the use of a uniform B-factor is justified by the high resolution of the data and the structural density of the pilus rod itself, which ensures a relatively uniform signal distribution throughout map. Sharpened maps are widely used in cryo-EM and are particularly valuable for water placement [26].

Compared to the unsharpened map, the sharpened one more clearly delineates the protein backbone and side-chain features (Fig. 3), enabling precise assignment of carbonyl groups orientation and the conformations of side chains. While the overall architecture of our pilus rod model closely resembles that of the previously published structure at 2.52 Å resolution (PDB: 8ptu, [13]), detailed comparison reveals that approximately two dozen residues per subunit adopt distinct conformations in our model. This is particularly important for the analysis of intersubunit interactions. For example, the sharpened map clearly resolves the hydroxyl group of Ser 91 in subunit F, allowing us to unambiguously position it at a hydrogen bond distance of 2.74 Å from the backbone nitrogen of Ala 25 in subunit H (Fig. 4). In contrast, the corresponding distances are longer in 8PTU model (3.25 Å, averaged across subunits) and in the recently reported enteroxigenic *E. coli* pilus structure (9IUG 3.29 Å [16]). This highlights the higher accuracy of our model.

**Figure 3.**
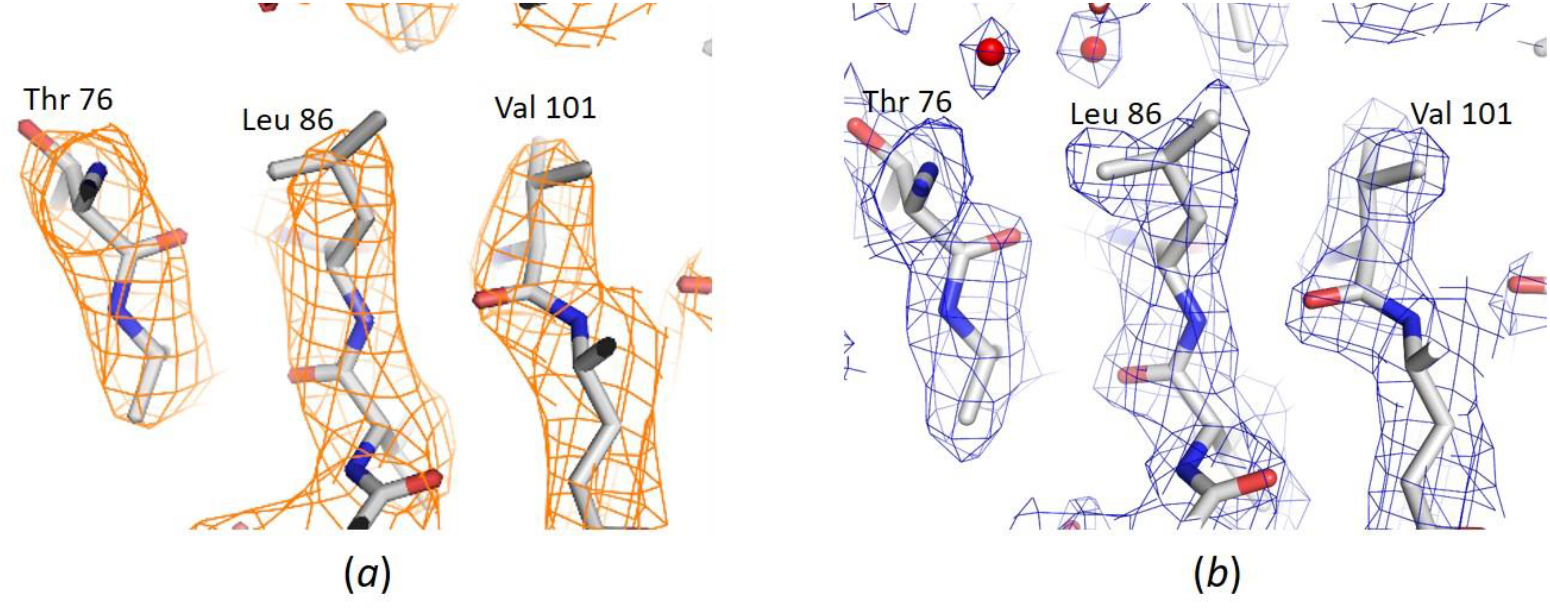
Comparison of the unsharpened (a) and sharpened (b) maps in the vicinity of Leu 86 of subunit H. (a) The unsharpened map is shown in orange at the contour level 0.18. (b) The sharpen map is shown in blue at the contour level 0.33.

**Figure 4.**
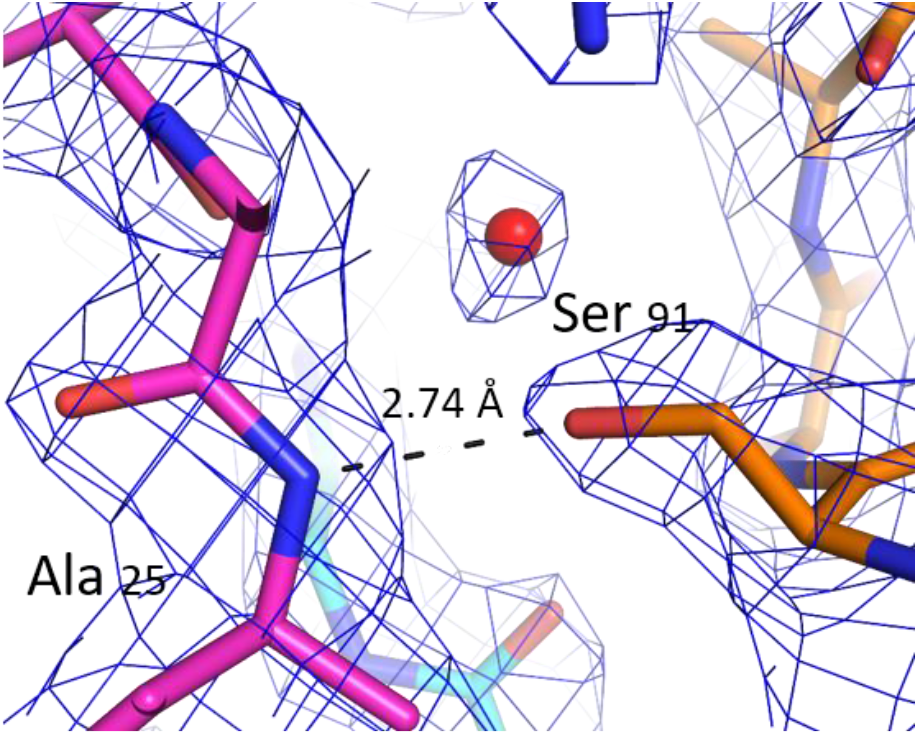
Close-up view of the interaction between Ser 91 of subunit F (shown in orange) and Ala 75 of subunit 25 (shown in magenta). The EM-map is shown at the contour level 0.34.

The most significant advantage of the sharpened map, however, is that it enabled us to construct a detailed water network around the pilus rod, which was not possible at lower resolution. Owing to the high concentration of hydrophobic residues in the core of each FimA subunit, water molecules are largely excluded from the interior (no more than two per subunit are present). Instead, water molecules are predominantly localized at the inner and outer surfaces of the rod and in intersubunit interfaces, where they contribute to subunit contacts. Notably, within the inner surface of the rod, each subunit contains a well-defined water network along β-strands 29–34 and 54– 57 (Fig. 1d), whose molecules exclusively interact with atoms of the same subunit.

The interactions between subunits within the pilus rod, previously described in earlier studies [4], reveal a complex organizational pattern. The incorporation of water molecules (this study) into the structural model provides new insights into the inter-subunit contacts. In addition to the well-established hydrogen bonding and hydrophobic interactions, two distinct roles of water molecules are observed: (i) individual water molecules that bridge residues from different subunits, and (ii) clusters of water molecules that form extended networks across subunits. The latter are typically associated with weaker and more diffuse density peaks. Both types of water-mediated interactions - bridging and network-forming - appear to contribute not only to the stabilization of the pilus architecture but also to its mechanical flexibility.

The analysis of water molecules in inter-subunit contact regions of our model revealed the following. Between neighboring subunits that differ in numbering by one unit (n and n+1), the contact is very tight. The β-strand of the n+1 subunit is inserted into a previous subunit n, and 22 hydrogen bonds with a length less than 3.2 Å with subunit n are formed (calculated by server PISA [27], which leaves not much space left for water networks. Only two water molecules connecting the side chains of neighboring subunits are observed. Notably one of these water molecules is surrounded by three subunits and is deep-buried inside the rod (Fig. 5a).

**Figure 5.**
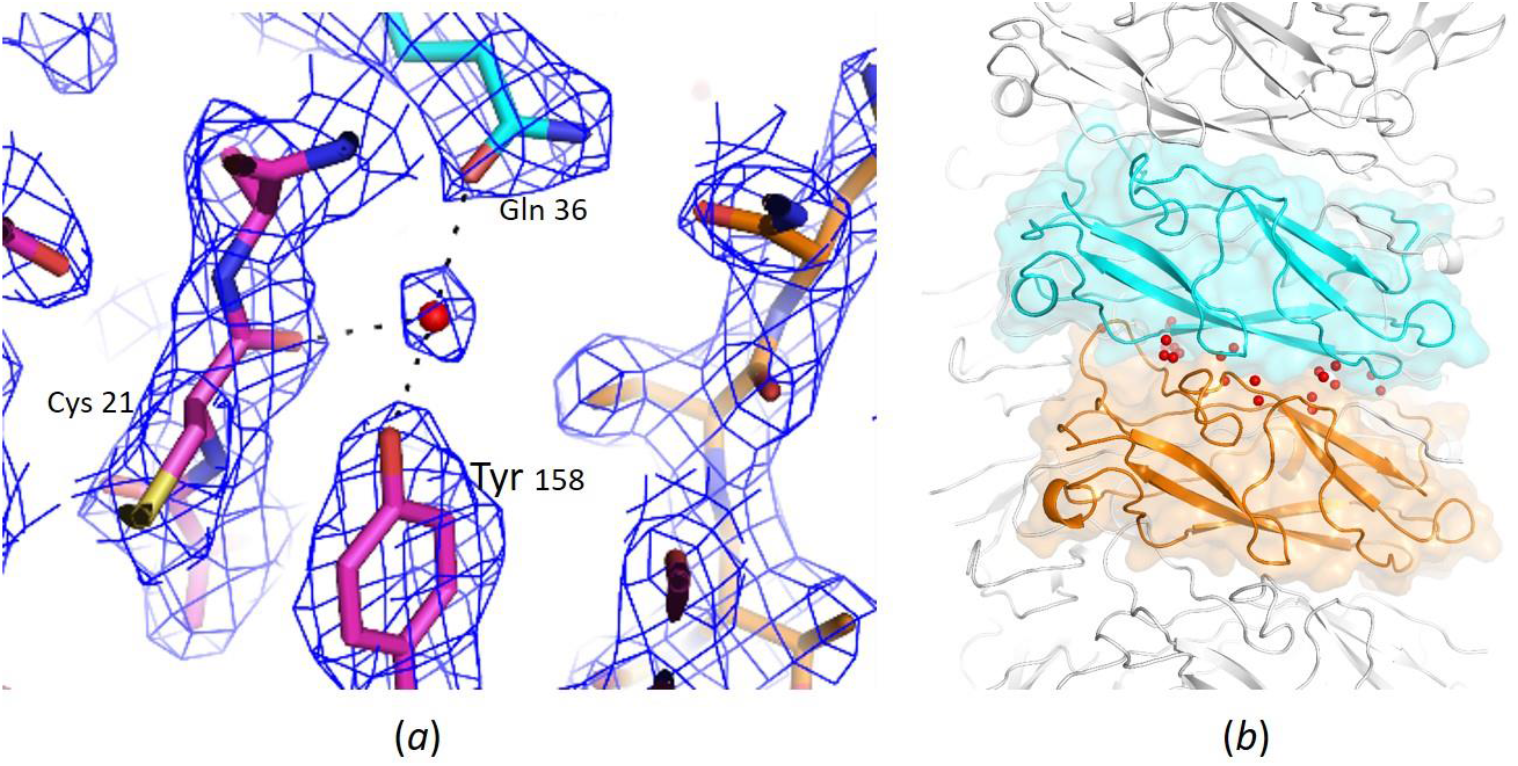
Water molecules participating in the contacts between subunits (a) One water molecule located at hydrogen bond distances from subunits H (shown in magenta) and I (shown in cyan). Subunit G is shown in orange. The EM-map is shown at the contour level 0.34. (b) The water network between subunits F (shown in orange) and I (shown in cyan).

The n and n+2 subunits have two spatially separated regions of contacts. In each of them, there is one hydrogen bond with a length of less than 3.2 Å between the subunits accompanied by one water molecule connecting the residues of these two subunits. In one of these regions, there is also a cluster of about five water molecules connecting the subunits.

In the region of contacts of n and n+3 subunits, there are five hydrogen bonds with a length of less than 3.2 Å, and instead of individual water molecules, there are a few clusters of water molecules forming networks between the subunits (Fig. 5b).

Interestingly, even between the n and n+4 subunits for which contacts are often not considered for type 1 pili as there are no hydrogen bonds, there are two water molecules connecting the residues of these two subunits (Figs. 6a,b).

**Figure 6.**
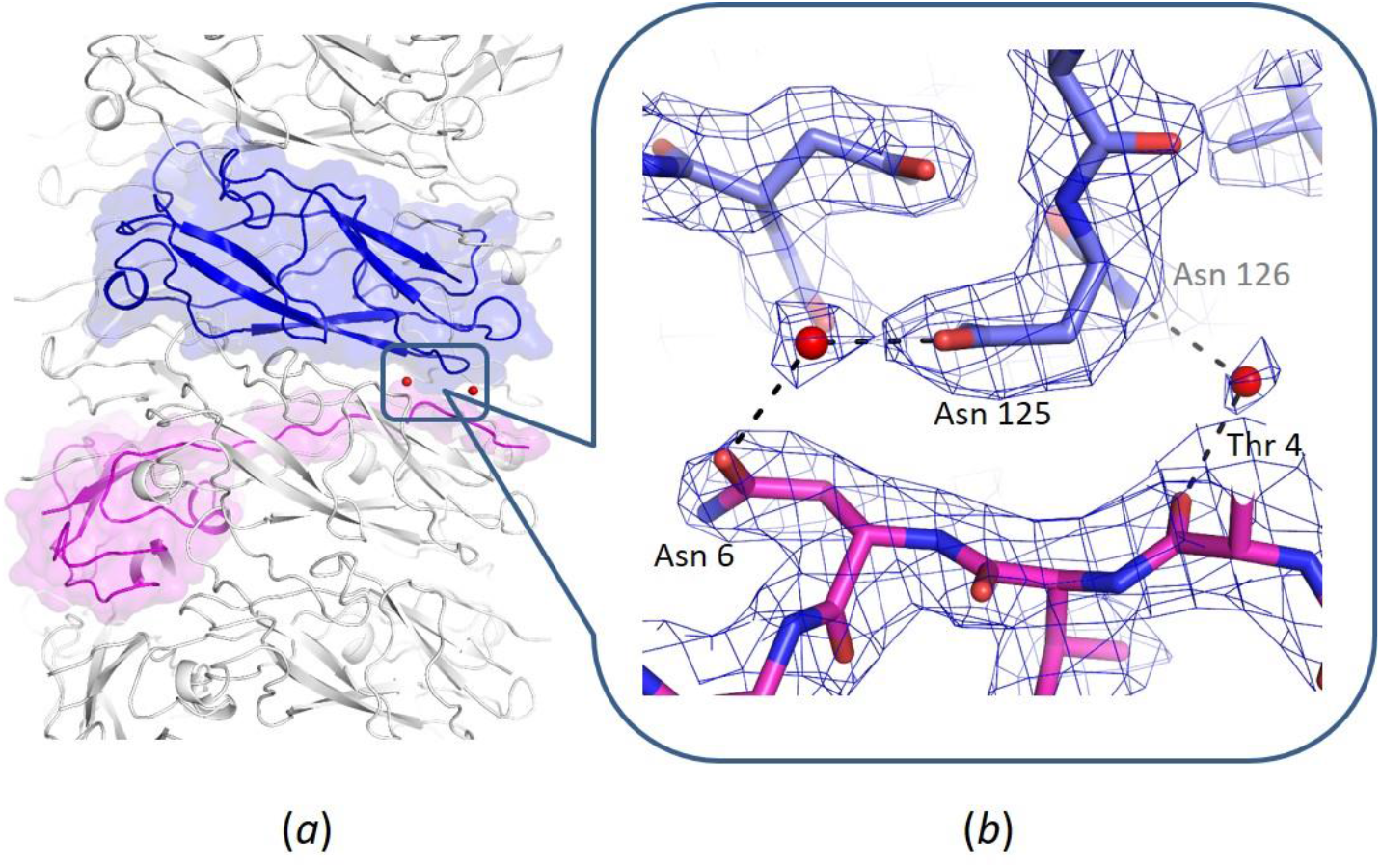
Water molecules connecting subunits E (shown in magenta) and I (shown in blue). (a) Subunits E and I in the pilus rod. (b) An enlarged view of the region around two water molecules connecting subunits E and I.

### 3.2. Evaluating the quality of the EM map and the model

A popular criterion in assessing the quality of maps and models in diffraction experiments is resolution, which refers to how well the details on the map are resolved. In electron microscopy, there are different concepts and definitions of the resolution and different approaches to how to calculate it.

To estimate the overall resolution of an EM reconstruction, it is a standard practice to calculate the Fourier Shell Correlation (FSC) between two independently refined half-maps. The spatial frequency at which the FSC curve falls to 0.143 is conventionally reported as the achieved resolution. This criterion reflects the highest spatial frequency at which the two half-maps contain consistent signal, thereby providing an estimate of the information limit of the reconstruction. It should be noted that cryoSPARC generates several FSC curves. In practice the FSC curve termed “corrected” is used to estimate the resolution, which often leads to the mismatch with the PDB resolution estimates. This curve is the result of applying a mask and is calculated using the high-resolution noise substitution procedure [28]. In the present study, cryoSPARC yields a resolution estimate of 2.5 Å, based on the “no mask” curve, consistent with the PDB estimate, and 2.09 Å from the “corrected” curve (Fig.1a), which appears to be overly optimistic. For comparison, the resolution estimate of 2.52 Å reported for the 8ptu structure, was obtained using a masked FSC curve [13]. Additional resolution estimates obtained using different programs are summarized in Table 2. One estimate was determined from the FSC-curve using the phenix.mtriage procedure [29], while another, termed d99, corresponds to the cutoff beyond which the Fourier map coefficients become negligibly small. The median local atomic image resolution, reflecting the local resolution of the experimental map around the centers of the atoms was calculated following the procedure described by Urzhumtsev and Lunin [30,31]. Together, these analyses demonstrate that the resolution in our study is improved (Table 2). Furthermore, the model Q-score, which quantifies the agreement between atomic coordinates and the experimental density [32], is also improved (Table 2). Since the Q-score is known to correlate with the map resolution [33], this improvement further supports the enhanced quality of our reconstruction.

**Table 2.**
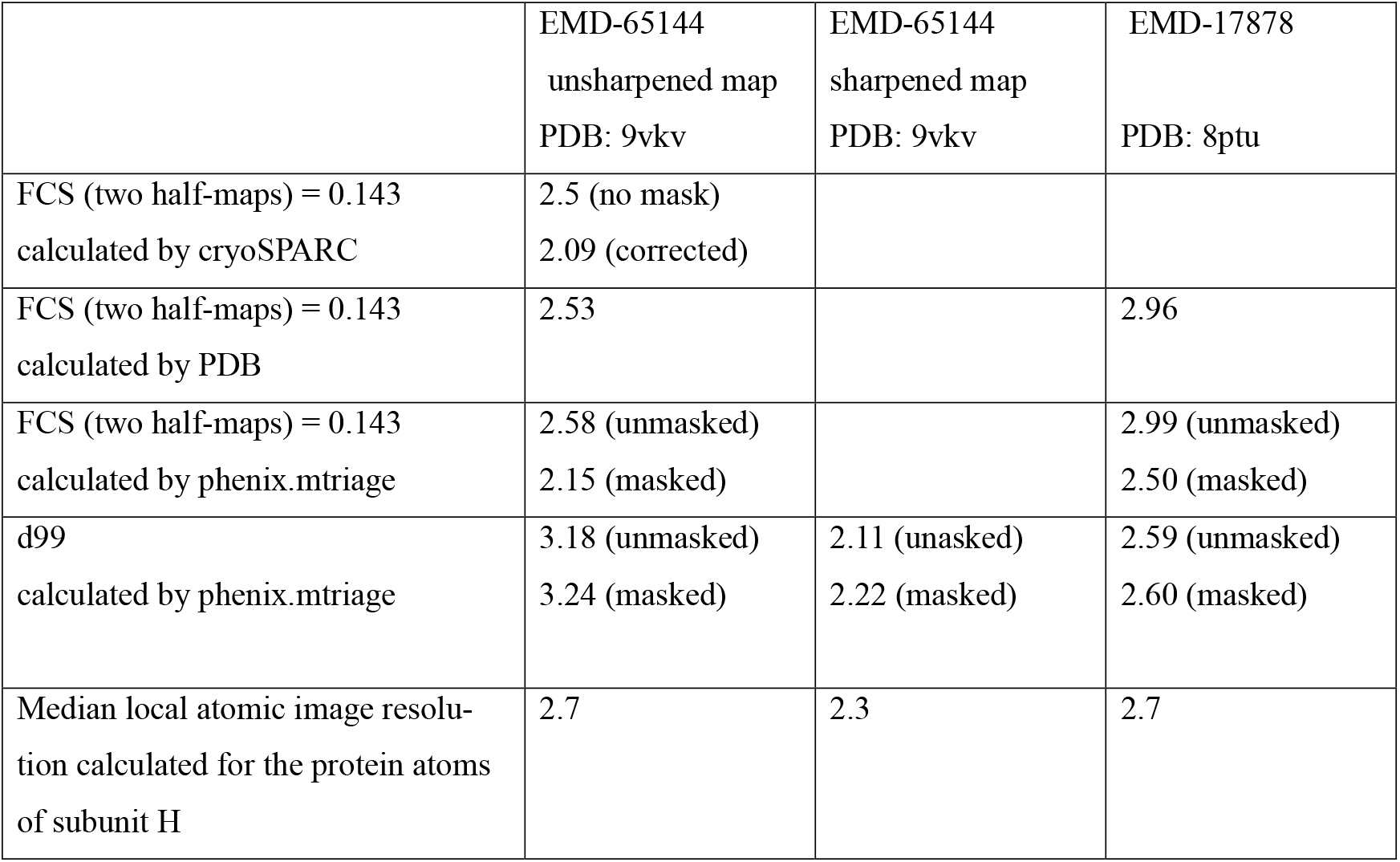
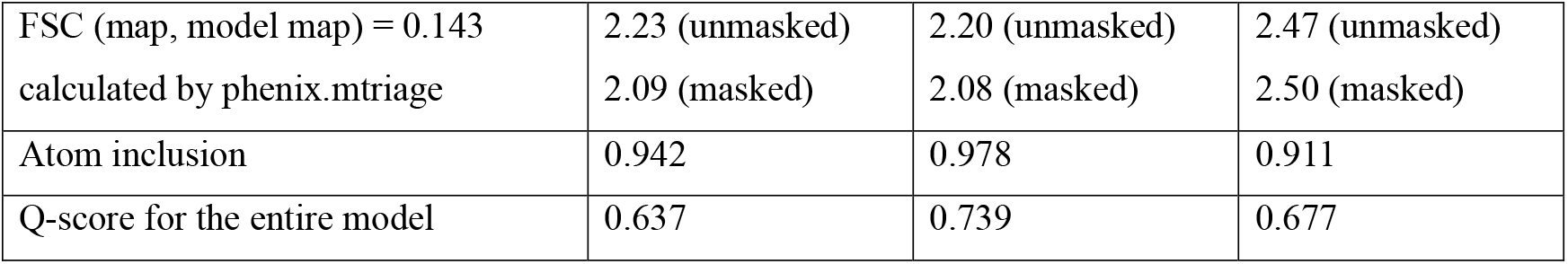
The resolution estimations of EM-map calculated using different approaches and programs and some indicators how models fit maps.

For the unsharpened map, the resolution estimates obtained without masking is equal to ∼2.5 Å according to a few different programs. The analysis of the sharpened map suggests an improved resolution, with realistic estimates falling within the range of 2.09-2.30 Å, depending on the program used.

### 3.3. Water network validation

The map sharpening procedure not only enhances the signal but also amplifies the noise, potentially introducing spurious density. For this reason, thorough validation of the water network model is essential. To reduce the risk of false-positive assignments when adding water molecules using phenix.douse, we set the attenuation scale factor (which defines whether a density peak can be considered as a possible water position) higher than the default value. In addition, all included water molecules satisfied basic geometric criteria: they were positioned at hydrogen bond distances from protein atoms and from other water molecules.

To further assess the reliability of the model, we compared our structure with a recently deposited pilus model of enterotoxigenic *E. coli*. The major subunit FimA in that model differs from FimA of our model by only 17 residues. We specifically examined whether water molecules in two models occupy equivalent positions.

As the first step, we selected only the “conserved” water molecules in our model, i.e., those that appear in equivalent positions across multiple subunits, rendering them the most reliably assigned. The superposition of five central subunits F, G, H, I, and J together with their waters followed by visual inspection in Coot revealed that of the 99 water molecules surrounding subunit H, 50 were also present in the other four subunits. Several of these conserved waters participate in intersubunit contacts, while others form a continuous network along two β-strands on the inner surface of the rod. (Fig. 1d).

Next, we compared these conserved water molecules with those in the model of the pili from enterotoxigenic *E. coli*. H subunits were superimposed and water molecules were considered equivalent if being no more 1 Å apart, to account for slight backbone misalignments and positional errors. It is remarkable that, 49 out of 50 conserved water molecules in our model coincided with the homologous pilus model of enterotoxigenic *E. coli*. This high degree of overlap indicates that nearly all reliably assigned water molecules are reproduced in independent experiments and in the homologous structures, supporting the validity of the water network in our model.

To identify structural differences we also compared the less reliable non-conserved water molecules. In subunit H, 27 waters were absent in the homologous structure. Of these, 11 water molecules were located near residues that differ between the two FimA sequences, with corresponding differences in positions of water molecules (Fig. 7), as expected. Among the remaining waters, 8 belonged to the second hydration shell (interacting only with other waters), while 6 were positioned in intersubunit regions. These observations suggest that sequence variation and local structural context influence water positions, and further support the accuracy of the water network defined in our model.

**Figure 7.**
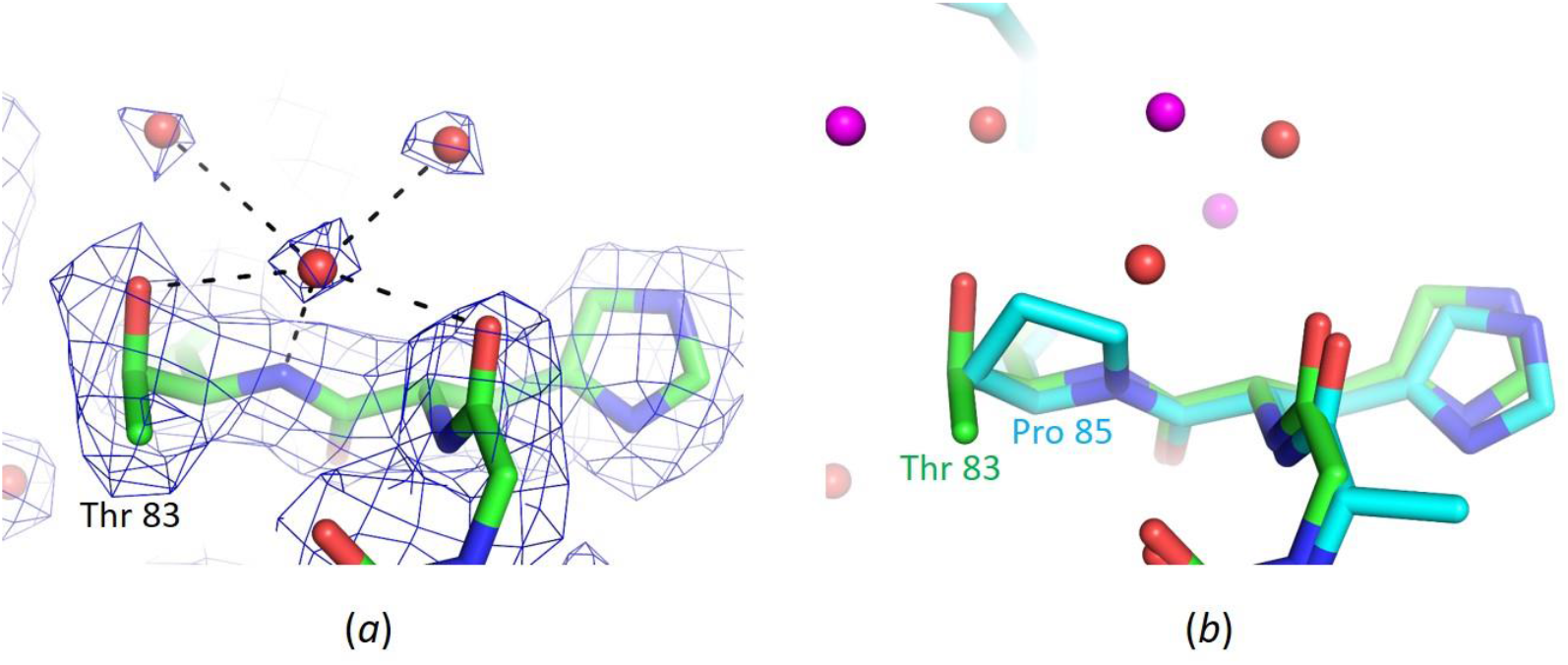
A comparison of water networks in the pilus models from UPEC K-12 and enterotoxigenic *E. coli*. (a) The model of pilus and water molecules (in red) in the vicinity of Thr 83 subunit H from UPEC K-12. The EM-map is shown at the contour level 0.22. (b) Superposition of the pilus rod from UPEC K-12 and the model of the pilus rod and water molecules (in magenta) from enterotoxigenic *E. coli*.

In summary, the high-resolution cryo-EM reconstruction of the type I pilus from UPEC enabled the reliable identification of a comprehensive water network within and between FimA subunits. Our analysis demonstrates that water molecules not only stabilize intersubunit interfaces through bridging interactions and extended networks but also contribute to the structural flexibility of the pilus rod. The strong agreement of conserved water positions with those observed in the homologous enterotoxigenic *E. coli* model further validates our assignments and underscores the functional significance of these hydration sites. At the same time, sequence-dependent differences in hydration highlight the adaptability of the water network to local structural environments. Taken together, these findings emphasize the essential role of water in maintaining both the stability and dynamic properties of pili, providing new insight into the molecular basis of bacterial adhesion and potential avenues for future therapeutic targeting.

